# Distinct Origins of Two-order Hierarchical Cognitive Abilities in Human Adults

**DOI:** 10.1101/2021.05.18.444677

**Authors:** Shaohan Jiang, Fanru Sun, Peijun Yuan, Yi Jiang, Xiaohong Wan

## Abstract

Human cognitive abilities are considerably diverse from basic perceptions to complex social behaviors. All human cognitive functions are principally categorized into a two-order hierarchy. Almost all of the first-order cognitive abilities investigated in behavioral genetics have been found to be dominantly heritable. However, the origins of the human second-order cognitive abilities in metacognition and mentalizing so far remain unclear. We here systematically compared the origins of the first-order and second-order cognitive abilities involved in the metacognition and mentalizing tasks using the classical twin paradigm on human adults. Our results demonstrated a double dissociation of the genetic and environmental contributions to the first-order and second-order cognitive abilities. All the first-order cognitive abilities involved in the metacognition and mentalizing tasks were dominantly heritable. In contrast, the shared environmental effects, rather than the genetic effects, had dominant contributions to the second-order cognitive abilities of metacognition and mentalizing in human adults. Hence, our findings suggest that human adults’ monitoring sensitivities in metacognition and mentalizing are profoundly sculpted by their social or cultural experiences, but less preconditioned by their biological nature.

## Introduction

Everyone has different talents. Some are good at motoric activities (*e.g*., playing soccer), while others are better in cognitive tasks (*e.g*., math). There is a long history in questing the origins of individual differences in human abilities, specifically in cognition (Galton, 1870; Plomin and Daniels, 2011). All human cognitive functions are principally categorized into a two-order hierarchy (Flavell, 1979; Vygotsky, 1978). The first-order cognitive functions refer to mental computations on external information from the physical world, while the second-order cognitive functions refer to mental computations on internal information from the mental world that is generated during the first-order cognitive processes. That is, the second-order cognitive processes monitor and manipulate the first-order cognitive processes (Nelson, 1990) (**Figure 1A**). Critically, the capabilities of the two-order hierarchical cognitive functions might be dissociable. For instance, the capabilities to monitor one’s own decisions (*i.e*., metacognition) are enormously varied even when the first-order performance accuracies in the decision-making tasks are controlled to be similar across individuals (Kunimoto et al., 2001; Fleming et al., 2010). We here test whether the genetic and environmental origins of individual differences in the two-order hierarchical cognitive abilities might be also distinct.

**Figure 1.**
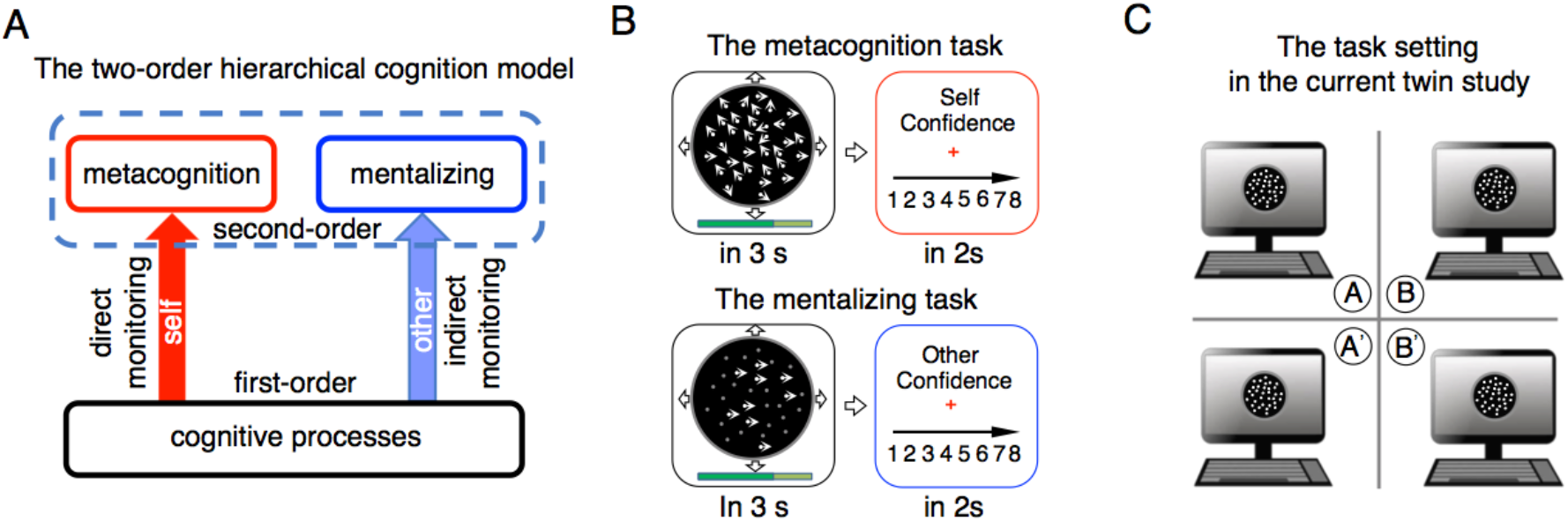
Experimental paradigm and task setting. (**A**) The two-order hierarchical cognition model. Both metacognition and mentalizing are the second-order cognitive processes in monitoring one’s own or the others’ first-order cognitive processes, respectively. (**B**) The experimental paradigms. In the metacognition task, the participant perceived and judged the net direction of random dot kinematogram (RDK) within 3 seconds (the elapsed time was shown as a progress bar under the stimulus), and then reported the confidence rating. The task difficulty was controlled by a staircase procedure prior to the experiment, so that the accuracy for each participant converged towards 50%. In the mentalizing task, the participant perceived noiseless moving-dots in that the originally randomly moving-dots in the metacognition task remained stationary. Concurrently, the participant also perceived the elapsed time that the partner used to make a choice. The participant then reported the estimate about the partner’s confidence rating. (**C**) On each occasion, two pairs of twins (either MZ or DZ) participated the experiment and conducted both the metacognition task and the mentalizing task twice. They were randomly paired with one another. When one participant was conducting the mentalizing task, another partner was concurrently conducting the metacognition task to estimate the former partner’s confidence from the RT information (the stimulus coherence was fixed). The four participants were physically separated from one another by board panels, while their computers were, if necessary, synchronized by network connection following the TCP/IP protocol.

The classical twin studies are the most promising and naturally available paradigm to investigate the biological and environmental origins of individual differences in human cognitive abilities (Neale and Cardon, 1992; Polderman et al., 2015). The monozygotic (MZ) twins have identical genes but the dizygotic (DZ) twins, on average, share a half of genes, while both the MZ and DZ twins are normally raised in the same family environments (Plomin and Daniels, 2011). Hence, a greater resemblance of a cognitive ability between the MZ than DZ twins reflects the existence of a genetic factor contribution, while a common resemblance between the MZ and DZ twins instead reflects the existence of shared family environmental contribution (Neale and Cardon, 1992). Over a century of research studies in behavior genetics using the classical twin paradigm have collectively demonstrated that almost all of the first-order human cognitive abilities are heritable (Turkheimer, 2000). About 50% of the variances of individual differences in twins’ first-order cognitive abilities are explainable by the common genes inherited from their same parents (Plomin and Deary, 2015; Polderman et al., 2015). Further, people with higher abilities on one cognitive task tend to also have higher abilities on the other tasks. The general cognitive ability [*i.e*., g or IQ (intelligence quotient)] captures the covariance across the first-order cognitive abilities and also well predicts the socioeconomic status (SES) and wellbeing (Deary et al., 2010; Strenze, 2007). In literature, general intelligence is reported to have 50~80% heritability (Plomin and Deary, 2015; Polderman et al., 2015).

However, general intelligence cannot similarly explain the second-order cognitive abilities, even though both significantly contribute to the cognitive performance in everyday life (Greven et al., 2009), but see also ref.(Sternberg, 1984). The second-order cognitive process in monitoring one’s own current first-order cognitive processes is referred to as metacognition (*e.g*., I believe how well I made the decision), while the corresponding process in monitoring the others’ cognitive processes are termed as mentalizing (*e.g*., I believe how well you believe you made the decision). Differing from metacognition that can directly introspect the internal information inside one’s own brain, mentalizing is unable to access the others’ internal information. Instead, it may rely on social cue associations, simulations of metacognitive experiences, or the theory of theory (*i.e*., theory of mind, ToM), to estimate the others’ mental states (Flavell, 2003). Hence, although the two second-order cognitive processes commonly involve in meta-representations of mental states, the underlying neural processes and mental state representations are different (Feurer et al., 2015).

Although the psychological and neurobiological mechanisms have been extensively explored (Frith and Frith, 2012; Fleming et al., 2012; Qiu et al., 2018), the biological and environmental origins of the second-order cognitive abilities of metacognition and mentalizing so far remain largely unclear. In light of the observations that both cognitive abilities even appear in preverbal infants (Goupil and Kouider, 2016; Onishi and Baillargeon, 2005), it is conceivable that the two second-order cognitive abilities might be also critically rooted in the genes, as similar as the first-order cognitive abilities. However, the acquired evidence from a few twin studies remains controversial on the genetic influences on the mentalizing abilities (Davies et al., 2016; Hughes et al., 2005; Knafo and Uzefovsky, 2013; Warrier et al., 2018). For the metacognitive abilities, to our knowledge, the twin studies are so far lacking, probably because the administration of the metacognition tasks is arduously demanding for a large sample size of participants, and reasonable confidence rating is unavailable for children.

## Results

To compare the genetic and shared family environmental contributions respectively to the first-order and second-order cognitive abilities, in the present study we recruited fifty-seven pairs of adult MZ twins and forty-eight pairs of adult DZ twins to participate both the metacognition and mentalizing tasks (**Table 1**). On each case, two pairs of twins (either MZ or DZ) came together and concurrently conducted both tasks (see **Materials and Methods**). In the metacognition task, the participant reported the confidence rating immediately after making a decision about the direction of random dot kinematogram (RDK) from four alternative options (**Figure 1B**, upper). The task difficulty was determined by the fraction of coherently moving dots (*i.e*., coherence), which was calibrated by a staircase procedure with the performance accuracy towards 50% for each participant prior to the experiment. In the mentalizing task, the participant concurrently observed the partner’s task performance on the RDK task and attributed the trial-by-trial confidence to the partner. Critically, to avoid evoking the participant’s own decision uncertainty on the RDK task, only those originally coherently moving dots remained moving, while the other originally randomly moving dots were stationary (**Figure 1B**, bottom). By virtue of this alternation, the RDK stimulus presented to the participant was noiseless, thereby, the participant could perceive the task difficulty, but would not have her/his own decision uncertainty as experienced in the metacognition task. Sequentially, the participant also perceived a progress bar representing the response time that the partner used to make the choice. The participant did not receive the information about the option choice and confidence rating reported by the partner. Each participant conducted the metacognition task and the mentalizing task twice. When the participant conducted the mentalizing task, the partner, either the sibling of the same twin (within-twin) or one of another pair of twin (between-twin), was concurrently conducting the metacognition task, and vice versa. The two pairs of twins were physically separated from one another, while the concurrent metacognition and mentalizing tasks were synchronized by network connection following the TCP/IP protocol via an Ethernet cable (**Figure 1C**).

**Table 1.** The MZ and DZ twins’ demographics and socioeconomic status (SES).

| | MZ | DZ | $\chi^2/T$ | *P* |
| --- | --- | --- | --- | --- |
| N (pairs) | 57 | 48 | — | — |
| % Girls | 48 | 58 | 1.0 | 0.31 |
| Age range | 20~30 | 20~29 | — | — |
| Mean age (SD) | 25.1 (2.4) | 23.4(2.8) | 1.5<sup>1</sup> | 0.14 |
| Self education in years<sup>2</sup> | 16 | 16 | 3.8 | 0.22 |
| Father's education<sup>2</sup> | 12 | 12 | 2.5 | 0.37 |
| Mother's education<sup>2</sup> | 12 | 12 | 1.4 | 0.26 |
| Self income per month<sup>2</sup> | 5~7k | 5~7k | 2.9 | 0.42 |
| Father's income<sup>2</sup> | 3~5k | 3~5k | 8.6 | 0.10 |
| Mother's income<sup>2</sup> | 3~5k | 3~5k | 8.8 | 0.13 |
| Father's occupation<sup>2</sup> | 3 | 3 | 3.8 | 0.43 |
| Mother's occupation<sup>2</sup> | 3 | 3 | 0.5 | 0.49 |

<sup>1</sup>T value; <sup>2</sup>the median.

There were no systematical differences between the MZ and DZ twins in either first-order or second-order behavioral measures related to the metacognition task (**Figure 1 - figure supplement 1**). As the participants performed the metacognition task twice, we could evaluate the reliability of these behavioral measures using the test-retest correlation. Except for the stimulus coherence that was the same across the two runs for each participant (**Figure 1 - figure supplement 1H**), the first-order behavioral measures in the RDK task (accuracy, median RT, RT variance) and even mean confidence (see discussion) had high reliability across the two runs (Pearson’s *r* = 0.68~0.95; Cronbach’s alpha: 0.78~0.95, **Figure 1 - figure supplement 1A-D**), while the second-order behavioral measures (RT-confidence correlation, gamma correlation between the confidence and correctness, raw AUC, and residual AUC) in metacognition had mediocre reliability (*r* = 0.50~0.72; Cronbach’s alpha: 0.50~0.80; **Figure 1 - figure supplement 1E-G**). These behavioral measures of the second-order metacognitive abilities had also high consistency (Cronbach’s alpha: 0.65~0.80 in the MZ and DZ twins, **Figure 1 - figure supplement 1I**).

We then calculated the intraclass correlation coefficients (ICCs) to measure the resemblance between the pairs within and cross the twins (see **Materials and Methods**). The within-twin MZ and DZ ICCs of the first-order and second-order behavioral measures were consistently higher than the corresponding cross-twin MZ and DZ ICCs (gray histograms, random pairs cross the MZ and DZ twins, respectively; **Figure 2 - figure supplement 1**), implicating substantial contributions from the same genes or the same family environments. Importantly, the ICCs of all the first-order behavioral measures within the MZ twins were consistently larger than those within the DZ twins (not significant in the RT variance), indicating considerable genetic factor contributions. In striking contrast, those of the second-order behavioral measures had no significant differences between the MZ and DZ twins, or some in the DZ twins were even numerically larger than the MZ twins (*e.g*., residual AUC), thereby indicating negligible genetic factor contributions.

To quantify the genetic and environmental contributions, we used maximum likelihood to compare all the potential structural equation models (SEMs) that decompose the covariance of each behavioral measure among the MZ and DZ twins into the different components associated with latent additive genetic factor (A), common shared family environmental (C), or non-additive (*i.e*., dominant) genetic factor (D), and non-shared family environmental factor (E, which also includes measurement errors), hereafter called the ACE or ADE model (**Equations 2** and **3** in **Materials and Methods**). We also took the correlated behavioral variables into account in the SEMs as the confounding variances (**Figure 1 - figure supplement 2**). The most parsimonious model with the smallest Akaike’s information criteria (AIC) was selected as the best model to account for individual variances in each behavioral measure.

As shown in **Table 2**, all the behavioral measures on the first-order cognitive abilities (coherence, median RT, RT variance, and mean confidence) were best accounted for by the AE or DE models (while the accuracy was controlled). The ratios of genetic contributions ranged from 0.36 to 0.41 in individual differences among the MZ and DZ twins. In striking contrast, all the behavioral measures on the second-order cognitive abilities (RT-confidence correlation, gamma correlation, raw AUC, and residual AUC) were best accounted for by the CE model. The ratios of shared family environmental contributions ranged from 0.17 to 0.40 in individual differences among the MZ and DZ twins. To further test the reliability of these model-comparison results, we repeated 100,000 times to randomly sample two-third of the MZ and DZ twins and made the same analysis procedure as described above (the bootstrapping procedure, see **Materials and Methods**). The dominant model by model comparison was still AE/DE for the first-order cognitive abilities, but CE for the second-order cognitive abilities (**Figure 2**). The dominance of the genetic factor or the shared family environmental factor in each behavioral phenotype in the full ACE model was the same as in the best-reduced model (**Table 2**). Hence, the dominant origin in each behavioral phenotype was not biased by the selected models (Hill et al., 2008).

**Figure 2.**
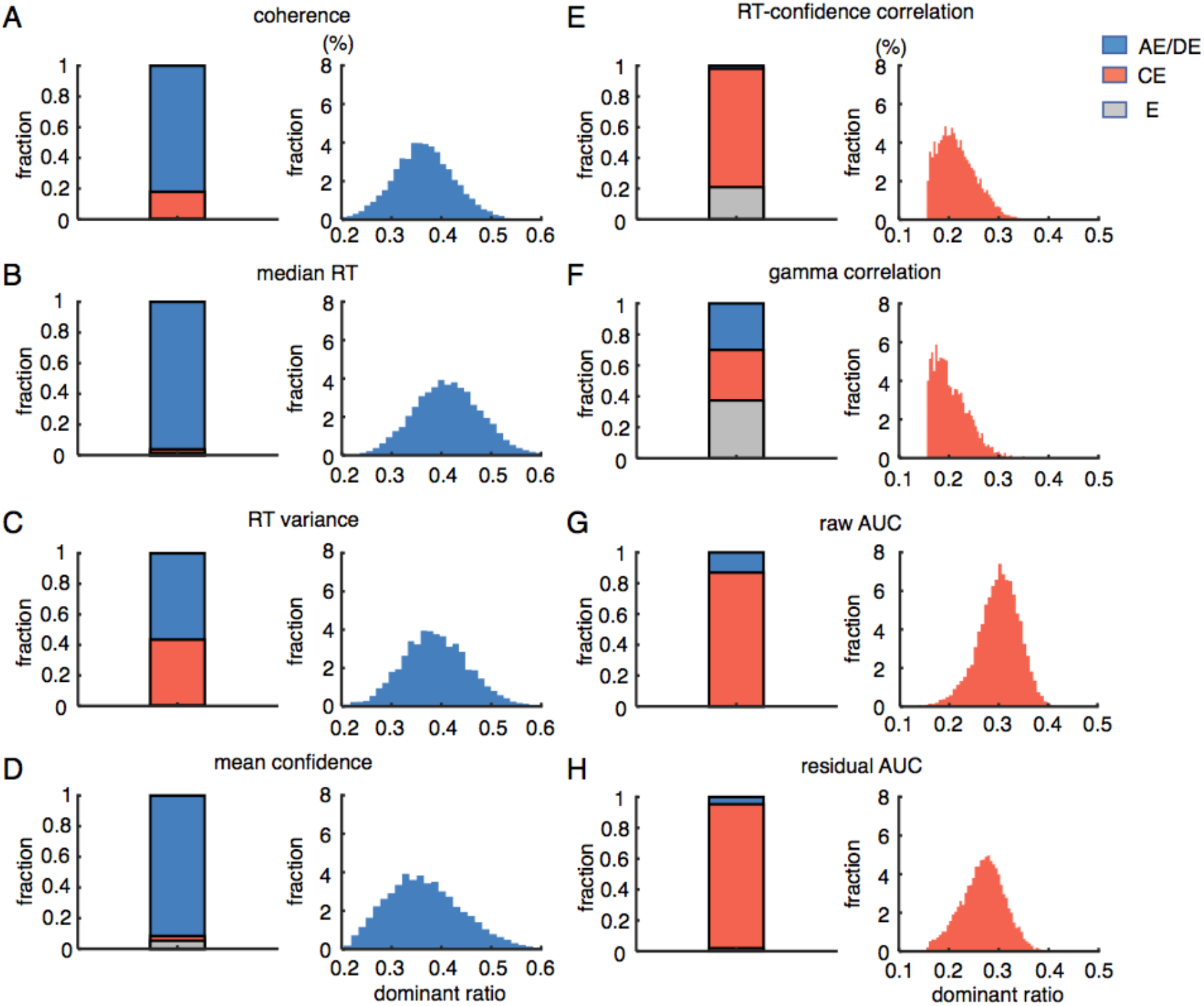
Distinct origins of first-order and second-order cognitive abilities in the metacognition task. (**A**) Coherence. (**B**) Median RT. (**C**) RT variance. (**D**) Mean confidence. (**E**) RT-confidence correlation. (**F**) Gamma correlation between the confidence and the correctness. (**G**) Raw AUC. (**H**) Residual AUC. The fraction ratios of the selected best models for the bootstrapping data and the dominant ratios of the shared genetic or environmental contributions in the behavioral measures of the metacognition task. The behavioral measures on the left column were the first-order cognitive abilities, while those on the right column were the second-order cognitive abilities.

**Table 2.** The estimated parameters of the behavioral measures in the metacognition task.

| phenotype | model | model fitting statistics |  |  | parameter estimates |  |  |
| --- | --- | --- | --- | --- | --- | --- | --- |
| | | $\Delta\chi^2$ | $P$ | $\Delta AIC^*$ | $a^2/d^2$ (95%C.I.) | $c^2$ (95%C.I.) | $e^2$ (95%C.I.) |
| coherence | AE | 0.001 | 0.97 | -2.04 | 0.36(0.25, 0.49) | 0 | 0.64(0.51, 0.75) |
|  | ACE | 11.96 | 0.06 | -0.04 | 0.36(0.16, 0.43) | 0(0, 0.11) | 0.64(0.57, 0.73) |
| median RT | DE | -2.7e-7 | 1.00 | -6.34 | 0.41(0.14, 0.63) | 0 | 0.59(0.37, 0.86) |
|  | ACE | 8.73 | 0.19 | -3.26 | 0.36(0.20, 0.44) | 0(0, 0.1) | 0.64(0.55, 0.75) |
| variance RT | AE | 0.2 | 0.66 | -12.12 | 0.36(0.17, 0.54) | 0 | 0.64(0.46, 0.83) |
|  | ACE | 1.68 | 0.95 | -10.32 | 0.21(0.14, 0.30) | 0.13(0.08, 0.19) | 0.66(0.54, 0.75) |
| confidence | DE | 0.01 | 0.92 | -10.2 | 0.36(0.14, 0.63) | 0 | 0.64(0.37, 0.86) |
|  | ACE | 4.1 | 0.67 | -7.9 | 0.33(0.16, 0.43) | 0(0, 0.1) | 0.67(0.57, 0.78) |
| RT-confidence | CE | 0.13 | 0.72 | -10.54 | 0 | 0.21(0.01,0.51) | 0.74(0.49, 0.99) |
|  | ACE | 8 | 0.24 | -8.54 | 0(0, 0.08) | 0.19(0.10,0.28) | 0.81(0.71, 1.00) |
| gamma corr | CE | 0.04 | 0.82 | -10.97 | 0 | 0.17(0.00,0.38) | 0.83(0.62, 1.00) |
|  | ACE | 3.03 | 0.8 | -8.97 | 0.09(0, 0.25) | 0.11(0.00,0.29) | 0.80(0.57, 1.00) |
| raw AUC | CE | 0.11 | 0.74 | -10.89 | 0 | 0.40(0.21,0.58) | 0.60(0.42, 0.79) |
|  | ACE | 3.11 | 0.79 | -8.89 | 0.10(0, 0.34) | 0.33(0.13,0.56) | 0.57(0.33, 0.83) |
| residual AUC | CE | -9.5e-6 | 1.00 | -10.25 | 0 | 0.22(0.03, 0.43) | 0.78(0.57, 0.97) |
|  | ACE | 3.75 | 0.71 | -8.25 | 0(0, 0.09) | 0.26(0.13, 0.36) | 0.74(0.65, 0.83) |

$\Delta AIC$ is the AIC difference of the listed model in reference with the saturated model.

In the mentalizing task, the participant estimated the partner’s decision confidence by observing the partner’s RTs, while the RDK stimulus coherence was stationary. The participant efficaciously used the RT information (the only available external information) to estimate the trial-by-trial decision confidence for either within-twin or between-twin partner. Thereby, the estimated confidence was highly correlated with the partner’s reported confidence (**Figure 3 - figure supplement 1A**), and could also predict the partner’s outcome (true or false). To measure the consistency between the estimated confidence and the partner’s outcome, we calculated the gamma correlation and AUC (**Figure 3 - figure supplement 1B and C**). There were no systematical differences between the behavioral measures in the four conditions of the mentalizing tasks. These behavioral measures of the mentalizing abilities had mediocre consistency with one another (Cronbach’s alpha: 0.4~0.7, **Figure 3 - figure supplement 1D**). However, after the RT-associated component was regressed out, the residual of the estimated confidence was not further correlated with the decision confidence reported by the partner, even from the same twin pair (**Figure 3 - figure supplement 1E**). The residual gamma correlation and the residual AUC calculated by the estimate residuals became also indifferent from the chance level (0 and 0.5, respectively; **Figure 3 - figure supplement 1F and G**). Nonetheless, the behavioral measures of the second-order mentalizing abilities had also mediocre consistency with one another (Cronbach’s alpha: 0.3~0.65, **Figure 3 - figure supplement 1H**). Hence, the participant relied on the RT-confidence association to make inferences on the others’ decision confidence.

Critically, most of the within-twin ICCs were significantly larger than the corresponding cross-twin ICCs (**Figure 3 - figure supplement 2**), indicating that the resemblance of the mentalizing ability was also specific for twins who shared the genes and family environment. Further, the participants seemed able to tell the partner from the same twin pair (within-twin) from the partner from another twin pair (between-twin), in that the respective ICCs were significantly differential (**Figure 3 - figure supplement 2**), even though they were physically separated from one another.

We thus separately compared the SEM models accounting for the within-twin and between-twin behavioral measures in the mentalizing tasks. For the within-twin behavioral measures, the SEM analyses also revealed that the individual differences of the RT-confidence association (RT weights), the gamma correlation, and the raw AUC among the MZ and DZ twins were dominantly accounted for by the AE/DE model (**Table 3**). The genetic contributions ranged from 0.36 to 0.52. In contrast, the individual differences of the residual AUC that was calculated after the RT-associated component was regressed out were best accounted for by the CE model (**Figure 3**). The shared family environmental contribution was about 0.19 (**Table 3**). For the between-twin behavioral measures, Similarly, the individual differences of the RT-confidence association and the gamma correlation were best accounted by the AE/DE model, but those of the raw AUC were best accounted by the CE model (**Figure 3**). The bootstrapping procedure further confirmed these results (**Figure 3**). The results in the full ACE model were as the same as in the best-reduced model (**Table 3**), with an exception of the AUC in between-twin mentalizing task. Thereby, the first-order cognitive ability of inference on the basis of the RT-confidence association in mentalizing was genetically influenced, but the second-order cognitive ability of mentalizing sensitivity beyond the RT-confidence association, particularly in the within-twin mentalizing task, was instead primarily influenced by the shared family environmental factor.

**Table 3.**
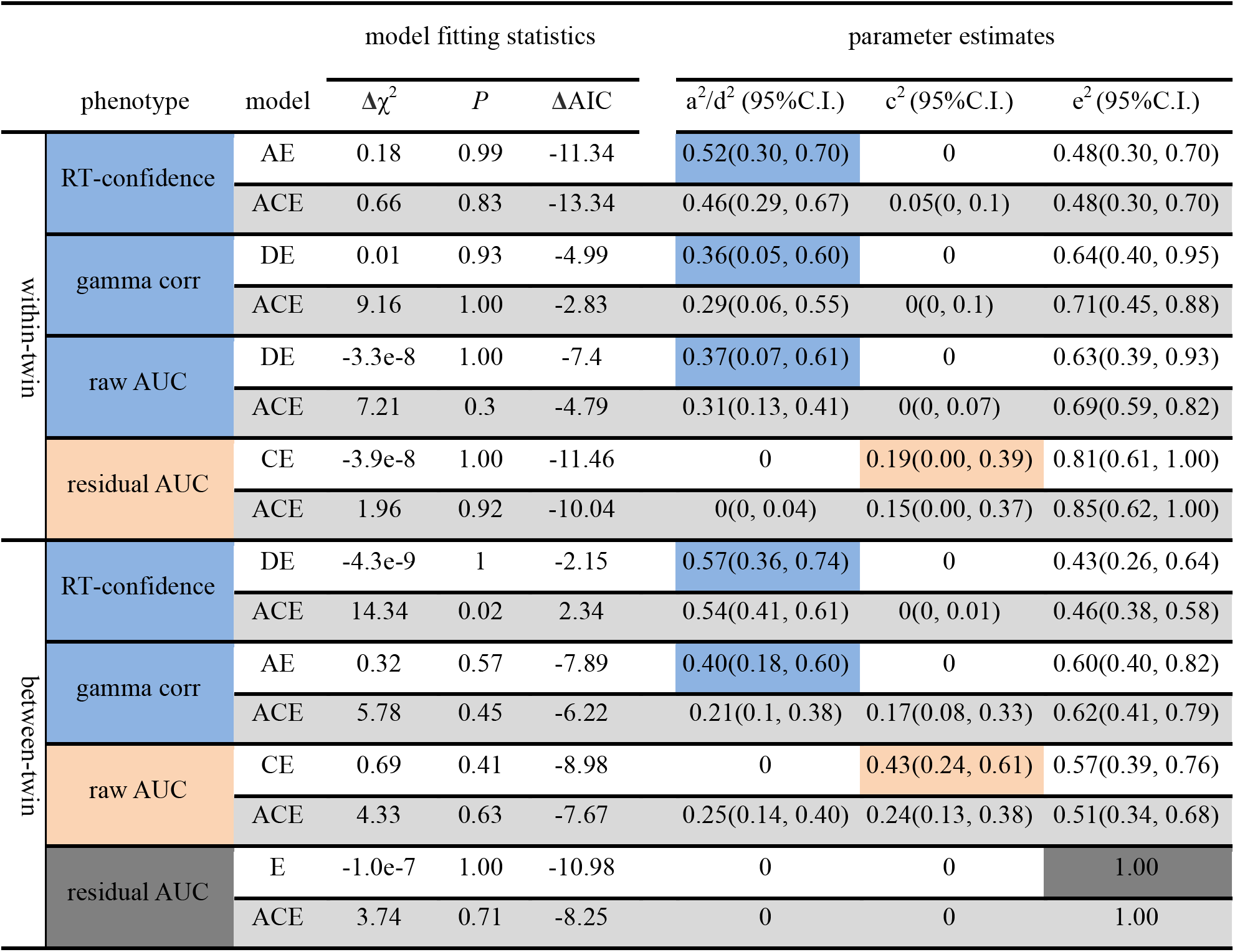
The estimated parameters of the behavioral measures in the mentalizing task.

**Figure 3.**
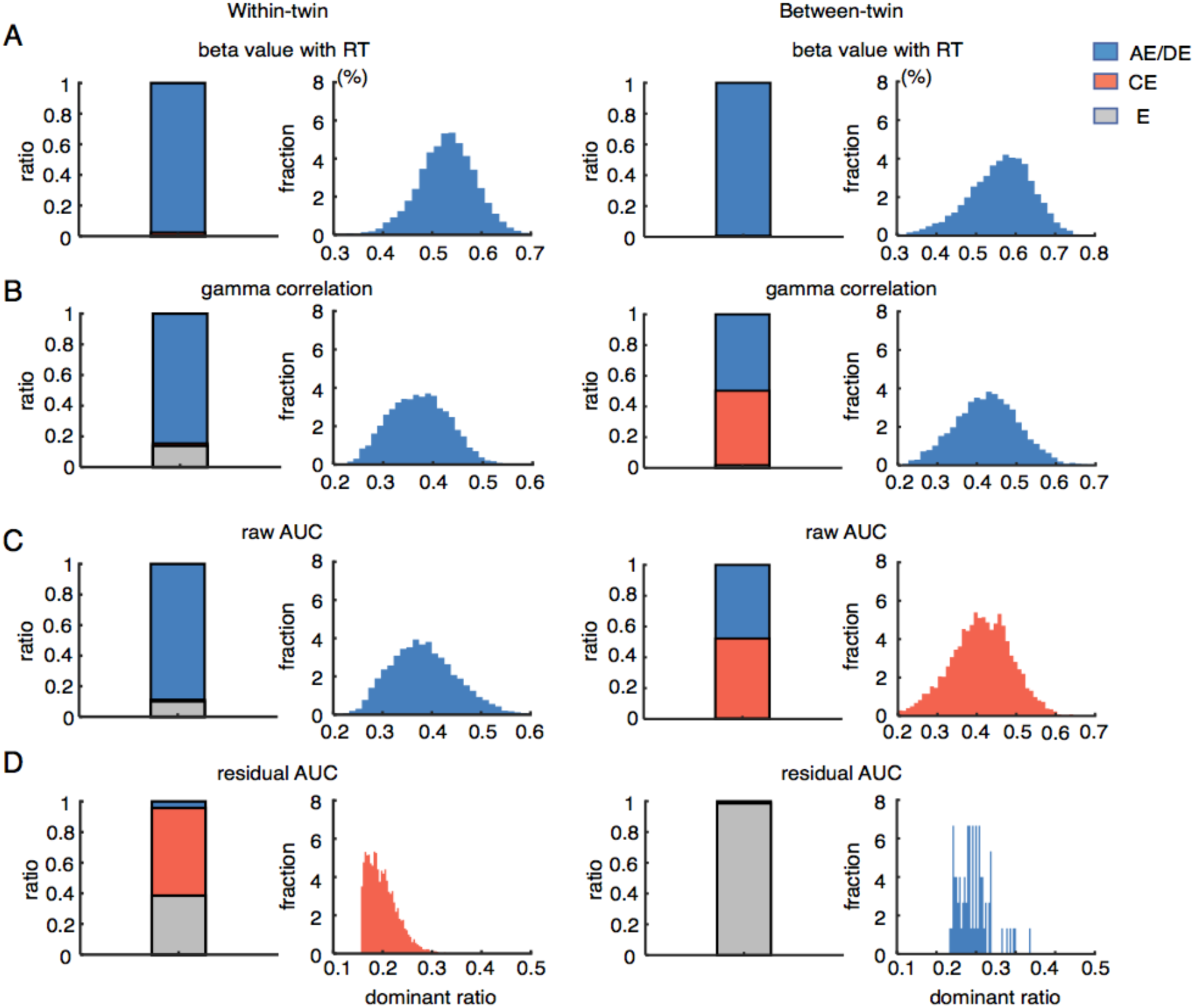
Distinct origins of first-order and second-order cognitive abilities in the mentalizing task. (**A**) Beta value of regression between the estimated confidence and the other’s RTs. (**B**) Gamma correlation between the estimated confidence and the other’s correctness. (**C**) Raw AUC. (**D**) The residual AUC after the RT-associated component of the estimated confidence was regressed out. The fraction ratios of the selected best models for the bootstrapping data and the dominant ratios of the shared genetic or environmental contributions in the behavioral measures of the mentalizing tasks. The behavioral measures on the left column were within the same twin (the participant estimated the confidence of the partner from the same pair of twin), while those on the right column were between the MZ and DZ twins (the participant estimated the confidence of the partner from another pair of twin).

However, unlike the assumptions that the common environmental factor and the non-additive (or dominant) genetic factor are alternately effective in the ACE and ADE models, the two factors might simultaneously affect the behavioral phenotypes (Keller and Coventry, 2005). In particular, ignorance of the non-additive or dominant genetic factor in the ACE model should cause the shared family environmental effect underestimated (Grayson, 1989). On the other hand, the potential confounding factors of gene-by-environment interactions, assortative mating, and sibling interactions might be misattributed to the shared family environmental factor, and thus cause it overestimated. To take accounts of these potential biases in the ACE and ADE models, we collapsed the additive and non-additive genetic factors as a single genetic factor (G) and used a *q*-GCE model (**Equations 5** and **6** in **Materials and Methods**) to assess the reliability of the genetic factor or the shared family environmental factor contributing to each behavioral phenotype by continuously changing the weight of the genetic factor (*q*) in DZ twins relative to MZ twins (Keller and Coventry, 2005). In a wide range of *q* values, the dominance of the genetic factor or the shared family environmental factor remained stable in contribution to each behavioral phenotype in both types of the tasks (**Figure 4**). Critically, the shared family environmental factor contributing to the residual AUC and the RT-confidence association in the metacognition task (**Figure 4A**), as well as the residual AUC in the within-twin mentalizing task (**Figure 4B**), even remained intact as the *q* values changed. Hence, the dominances of the genetic factor for the first-order cognitive abilities and those of the shared family environmental factor for the second-order cognitive abilities should be valid in a considerable variety of model space.

**Figure 4.**
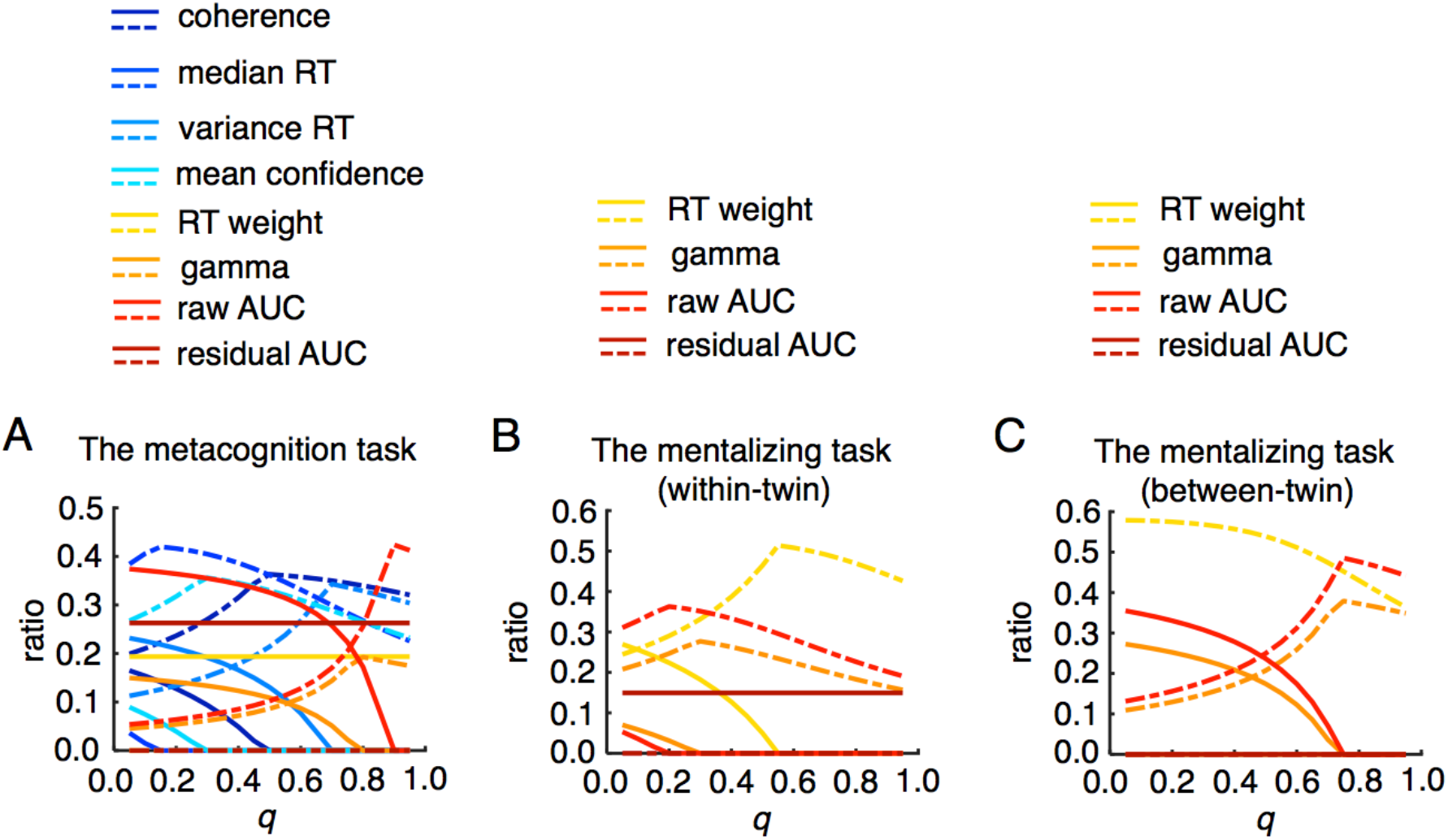
The contributions of the genetic factor and the shared family environmental factor changed with the *q* values in the *q*-GCE model. (**A**) The contribution ratios of the genetic factor (broken lines) and the shared family environmental factor (solid lines) changed with the *q* values in the metacognition task. The contribution ratio of the genetic factor was always larger than that of the shared family environmental factor in each behavioral measure of the first-order cognitive abilities in the entire range of *q* values [0.05, 0.95]. Instead, the contribution ratio of the shared family environmental factor was larger than that of the genetic factor in the behavioral measures of the second-order cognitive abilities in a larger range of *q* values. (**B**) The contribution ratios of the genetic factor (broken lines) and the shared family environmental factor (solid lines) changed with the *q* values in the within-twin mentalizing task. (**C**) The contribution ratios of the genetic factor (broken lines) and the shared family environmental factor (solid lines) changed with the *q* values in the between-twin mentalizing task.

A pitfall in the present study was that the sample size of the MZ and DZ twins was relatively small. This may raise concern about the robustness about the results. To assess the robustness of the results, we made post-hoc power analyses on the simulated data with the same covariance in the MZ and DZ twins obtained from the conventional ACE model for each behavioral phenotype using the OpenMx module (**Materials and Methods**). Although the post-hoc power with the sample size used in the present study was mostly low, even under 0.2 (**Figure 4 - figure supplement 1A**), the selectivity of contribution between the genetic factor and the shared family environmental factor 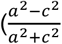, where a^2^ is the contribution of the genetic factor and c^2^ is the contribution of the shared family environmental factor) under the current sample size from the simulated data had systematic biases between the first-order and second-order cognitive abilities, consisting with the dominant factors revealed by the empirical data (**Figure 4 - figure supplement 2B**). Hence, future twin studies with a larger sample size are deserved to confirm the current findings.

## Discussion

Given that the genetic effect has virtually dominant influences on almost all the cognitive abilities and behavioral traits investigated in the literature of the classical twin studies (Polderman et al., 2015; Turkheimer, 2000), it should be much expectable of our observations that all of the first-order cognitive abilities examined in the current study were dominantly heritable. However, it is exceptional that the genetic factor had less influence on the second-order cognitive abilities of metacognition and mentalizing in human adults. On the contrary, the shared family environmental factor had considerable contributions to these cognitive abilities. These results should be not biased by the models used in the analyses and the noises in the current data, and thus should be qualitatively reliable. First, our results replicated the previous finding of no significant genetic influences on the mentalizing abilities (*i.e*., ToM, the false-belief tasks) with a quite large sample size of children at five years old (1,104 pairs of twins, in Goupil and Kouider, 2016), as well as in the recent studies (Warrier et al., 2018), even though the mentalizing tasks were different from the present study. Second, if the genetic effect would stably exist in a cognitive ability, the heritability in adults should be greater, rather than less, than that in children, due to potentially positive gene-environment interactions along the cognitive development (Johnson et al., 2009; Tucker-Drob et al., 2013). The genetic effects on the cognitive abilities are amplified under adequate environmental opportunities that are largely influenced by SES. There were no systematic differences in SES between the MZ and DZ twins who participated the experiment in the present study (**Table 1**). Third, identification of the shared family environmental effect usually should need a greater power than identification of the genetic effect, as the shared family environmental effect is usually more sensitive to noises (Hughes et al., 2005; Turkheimer, 2000), and underestimated by the conventional ACE model (Grayson, 1989; Keller and Coventry, 2005). Fourth, the bootstrapping results and the tests in a larger range of variational *q*-GCE models indicated that the observed effects were quite stable in the dataset. Lastly but most importantly, the genetic and environmental influences on the cognitive abilities in association with the metacognition and mentalizing tasks were clear-cut along the boundary of the two-order hierarchy: all of the first-order cognitive abilities were dominantly influenced by the genetic factor, but all of the second-order cognitive abilities were conversely influenced by the shared family environmental factor. The odds of the results by randomness should be very low [p = (1/2)^8^ = 0.0039 in the metacognition task]. This double dissociation of the dominant influences from the genetic factor and the shared family environmental factor on the first-order and second-order cognitive abilities profoundly implicates nature and nurture of human cognitive abilities.

In terms of the cognitive abilities associated with metacognition, the metacognition bias (mean confidence) was heritable (Plomin et al., 1992), but the metacognition sensitivity was not. Notably, the two behavioral measures were operationally independent. The people often have the trend to overestimate their performance (the Dunning-Kruger effect, Kruger and Dunning, 1999), even immediately after explicit learning from the actual outcomes (Kahneman and Lovallo, 1993). On the contrary, the metacognition sensitivity in discriminating the confidence levels with reference to the actual outcome is improvable with extensive learning from one’s own experiences and the others’ instructions. In fact, the explicit metacognition competence of reporting the confidence is not yet available at the early age of infants, but is gradually developed to maturation later in children around 8-9 years old, close to the ability in adults (Flavell, 1999). On the contrary, implicit metacognition competence of automatic monitoring decision uncertainty is ready even for preverbal infants (Goupil and Kouider, 2016).

On the other hand, in terms of the mentalizing abilities (*i.e*., cognitive empathy, ToM), the inference on the basis of the association with the external cue (*i.e*., RT) was heritable, but the intrinsic mentalizing sensitivity was not. The two abilities were also dissociated. The former was a first-order cognitive ability, but played a major role in mindreading (Kliemann and Adolphs, 2018; Lurz, 2011), in that the latter was not further able to predict the other’s performance after the cue-associated component was regressed out. However, these cue-independent residuals were not simply noises, but were, at least partially, generated from the neural activities in the social brain regions (*e.g*., the dorsomedial prefrontal cortex, Jiang et al. submitted). The neural computations in these brain regions should veridically reflect the mentalizing ability to estimate the inaccessible intrinsic mental states.

Any cognitive function is built-in on the intrinsic structures of the human brain that are predetermined by the genetic substances. During cognitive development, each functional capacity is also sculpted by the experiences during interactions with the physical and mental worlds. In most cases, the experiences should enhance the gene-driven diversities through the positive gene-environment interactions, but in some cases the experiences might dilute the genetic imprints and then become dominated.

The first-order cognitive functions are involved in processing of the external information from the physical world that is similar to all individuals and across generations. Accordingly, the anatomical structures in the brain areas associated with the first-order cognitive processes are both evolutionally and developmentally stable (Hill et al., 2010; Kaas, 2006). The intrinsic functional connectivity across these brain regions remains also less diverse during protracting maturation from the infants to adults within and across individuals (Gao et al., 2014; Mueller et al., 2013). In contrast, the second-order cognitive functions of metacognition and mentalizing specifically deal with the internal information in the mental world that is abstracted from the first-order cognitive processes. These functions are full of subjectivity, volatility, and idiosyncrasy. Metacognition is associated with the frontoparietal control network (Qiu et al., 2018), while mentalizing is associated with the social brain network (Frith and Frith, 2012). The anatomical structures in the two brain networks are instead much evolutionally and developmentally expanded (Hill et al., 2010; Kaas, 2006). The intrinsic functional connectivity across these brain areas remains also greatly diverse during protracting maturation from the infants to adults within and across individuals (Gao et al., 2014; Mueller et al., 2013). Although both the anatomical structures and intrinsic functional connectivity should be genetically influenced (Colclough et al., 2017; Jansen et al., 2015), the monitoring sensitivities in both metacognition and mentalizing are subject to the trial-by-trial neural activities in these brain areas matching the actual outcomes (*i.e*., AUC). Obviously, high predictability of the outcomes by metacognition and mentalizing needs arduous learning and experiences. The metacognition and mentalizing competencies become gradually matured until adolescents. During cognitive development, one important influence factor is presumed to be instructions that are received from the parents and teachers (Baker, 1994; Vygotsky, 1978). The cultural learning (Atzil et al., 2018; Heyes et al., 2020; Heyes and Frith, 2014; Vygotsky, 1978), for instance, the maternal talks about the children’s minds (Taumoepeau and Ruffman, 2008), may play important and unique roles in guiding children to understand their own and the caregiver’s mental states. Hence, the shared social environmental factors, such as parents’ caring, and so forth, should significantly shape the twins’ mental state representations in metacognition and mentalizing.

In conclusion, our results in the present study demonstrated that the dissociated first-order and second-order cognitive functions have distinct origins: the first-order cognitive abilities dominantly have genetic origins, but the second-order cognitive abilities dominantly have social origins. This finding implicates that the second-order cognitive abilities (monitoring sensitivities) in metacognition and mentalizing, unlike the first-order cognitive abilities, should be considerably malleable by social or cultural experiences during cognitive development, but less constrained by the biological nature. Thus, it is theoretically potential to develop efficacious education interventions in better fostering cognitive development in metacognition and mentalizing, and in better relieving the metacognitive and mentalizing deficits in psychiatric patients.

## Materials and Methods

### Participants

Two hundred and twenty-six right-handed healthy participants (sixty-five pairs of MZ twins and fifty-five pairs of DZ twins with the same sex, 24.2 ± 2.6 years old, one hundred and eighteen females) participated the present study (see **Table 1** for their demographics and socioeconomic status). The twins were recruited from a twin database (BeTwiSt) maintained by the Institute of Psychology, Chinese Academy of Sciences (IPCAS). Informed consent was obtained from each participant in accordance with a protocol approved by the institutional review board of IPCAS.

### Experimental procedures

#### Stimuli

In the metacognition task, stimuli appeared in an aperture with a radius of three degrees (visual angle); three hundred white dots (radius: 0.08 degrees, density: 2.0%) moved in different directions with a speed of 8.0 degrees/second on a black background. The lifetime of each dot lasted for three frames. A fraction of the dots moved toward the same direction (one of the four directions: left, down, right, and up), but the others moved toward different random directions. This was so-called a random-dot-kinematogram (RDK) stimulus (Kiani and Shadlen, 2009). In the mentalizing task, different from the stimuli used in the metacognition task, only the originally coherently moving dots remained moving, but those randomly moving dots remained stationary. This alternation of stimulus presentation could inform the participant about the stimulus difficulty (coherence), but would not evoke the participant’s own decision uncertainty about the net moving direction of the dots.

#### Metacognition task

This task was to assess the participant’s metacognitive ability in monitoring his/her own decision confidence or uncertainty. In each trial, the participant perceived the RDK stimulus surrounded by four direction arrows (left, down, right, and up) and made a judgment on the net moving direction in 3 seconds [the elapsed response time (RT) since the stimulus onset was simultaneously illustrated on the bottom of the screen. The green, yellow, and red color indicated that the elapsed time was within 1, 2, and 3 seconds, respectively], and subsequently reported the confidence rating with regard to his/her belief about the probability that the decision was correct in 2 seconds. The confidence ratings were on a scale from 1 to 8, where 1 indicated that the participant believed that his/her decision was completely wrong and 8 indicated that his/her decision was completely correct. No feedback about the correct answer was given (**Figure 1B**). The RDM coherence was constant in all trials and was set to match the task difficulty between different participants, such that the accuracy rate for each participant was about 0.5. The actual accuracy had some variations across the participants. The mean actual accuracy across the participants in the MZ and DZ twins was 0.47 and 0.48, respectively (**Figure 1 - figure supplement 1B**).

#### Mentalizing task

This task was contrastingly to assess the participant’s mentalizing ability in monitoring the other’s decision confidence or uncertainty, an internal mental state that was inaccessible, while the participant concurrently observed the partner’s behavioral performance on the metacognition task (**Figure 1B**). To preclude the participant from evoking his/her own decision uncertainty with the same RDK stimulus that the partner perceived, the participant actually viewed a different stimulus, in which only the originally coherently moving dots remained moving but the others were stationary. Concurrently, the participant also perceived the elapsed time that the partner was used up for making a choice. The partner’s choice and confidence rating were not revealed to the participant. Hence, for the participant, the only available information about the partner’s task performance was the RT, while the stimulus coherence remained constant. The participant subsequently reported his/her estimate about the partner’s confidence rating in 2 seconds.

#### Training

On the basis of our experience, performance on the metacognition task could become more stable after participants received sufficient practice. Therefore, each participant was trained for forty minutes prior to performing the two tasks. The RDK stimulus coherence (task difficulty) was adaptively adjusted trial-by-trial with a staircase procedure, such that the RDK stimulus coherence was upgraded one level after two continuous correct trials, downgraded one level after two continuous erroneous trials, and otherwise remained unchanged (Levitt, 1971). The RDK stimulus coherence started from 50%, and thereafter reduced and later remained at a stable level throughout the procedure. The performance accuracy rate became close to 0.5 at the end of practice. Because their stimulus coherences at the accuracy of 0.5 were larger than 20% (indicating that their perceptions on the RDK stimuli were not normal, mean ± S.D.: 11.8 ± 0.6 in the left participants), eight pairs of the MZ twins and seven pairs of the DZ twins were then excluded.

#### Experiment

On each case, two pairs of twins (either MZ or DZ) came together to participate the experiment for about one and half an hour. Each of them respectively conducted the metacognition task and the mentalizing task twice. When the participant conducted the mentalizing task, the partner, either the sibling of the same pair of twin (*i.e*., within-twin) or one of another pair of twin (*i.e*., between-twin), was concurrently conducting the metacognition task, and vice versa. The participants were physically separated from one another by board panels (**Figure 1C**). The concurrent metacognition and mentalizing tasks were synchronized by network connection following the TCP/IP protocol via an Ethernet cable. Each task was consisted of ninety trials and lasted ten minutes.

#### Socioeconomic status (SES)

To evaluate SES for each participant, we collected each participant’s education levels and his/her parents’ education levels (1. Middle school; 2. High school; 3. College; 4. Postgraduate); each participant’s income per month and his/her parents’ income per month (1. < 1,000 Yuan; 2. 1,000~3,000 Yuan; 3. 3,000~5,000 Yuan; 4. 5,000~7,000 Yuan; 5. 7,000~10,000 Yuan; 6. >10,000 Yuan); as well as his/her parents’ occupations (1. unemployed and retired; 2. partly skilled; 3. skilled; 4. professional and technical; 5. governmental and managerial). The medians of these SES measures were listed in **Table 1**.

### Data processing procedures

#### Statistical tests

Statistical analyses of data were performed with the Matlab statistical toolbox (Matlab2019, Mathworks, USA). Two-sample t-test or z-test was used to test whether each behavioral measure was significantly different between the MZ and DZ twins, and between within-twins and cross-twins. Two-sample K-S tests were used to compare the populations by bootstrapping with α = 0.05.

#### Individual metacognition and mentalizing abilities

A nonparametric approach was employed to assess each participant’s metacognition and mentalizing abilities (sensitivities) in monitoring decision-making, respectively. A receiver operating characteristic (ROC) curve was constructed by characterizing the correct probabilities with different confidence ratings as judgment criteria. The area under the ROC curve (AUC) was calculated to represent how precisely the participant was able to predict decision outcomes (Fleming et al., 2010). In other words, a larger AUC indicates the participant’s metacognition or mentalizing sensitivity is higher.

#### Behavioral measures

In the metacognition task, we separately measured each participant’s first-order and second-order cognitive abilities. The behavioral measures of the first-order cognitive abilities were constituted of stimulus coherence or accuracy, median RT, RT variance, and the mean confidence, while the behavioral measures of the second-order cognitive abilities were constituted of the RT-confidence correlation, the gamma correlation between the confidence and the correctness, the raw AUC, and the residual AUC after the RT-associated component was regressed out from the reported confidence. In the mentalizing task, as the participants used the RT information to estimate the partner’s decision confidence. We then calculated the beta value of the estimated confidence regressed with the RTs, the gamma correlation between the estimated confidence and the correctness, the raw AUC between the estimated confidence and the correctness. As much expected, these behavioral measures were mainly influenced by the association with the RTs. Thereby, these behavioral measures were considered as the first-order cognitive abilities. We then regressed out the RT-associated component from the estimated confidence, and recalculated the residual AUC as the second-order mentalizing ability.

#### Intraclass correlation coefficients (ICCs)

To measure the familial resemblance of each behavioral measure between the pair of twin, we calculated the ICCs separately for the MZ and DZ twins using one-way analysis of variance (ANOVA) random-effect model to account for the source of variance among the pairs of twins, regardless of the ordering on each twin (McGraw and Wong, 1996). That is

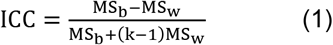

where MS_R_ and MS_W_ are mean square of variances between and within twins. *k* is specific to 2 in the twin study. Hence, if the variances between twins are considerably larger than those within twins, then the ICC value should be larger, and vice versa. We collapsed the data of the twice metacognition tasks, but separately dealt with the data of the mentalizing task with the sibling of the same pair of twin as the observational partner and the data of the mentalizing task with one of another twin as the observational partner, as the two datasets were inhomogeneous (**Figure 3**).

#### Modeling

To quantify the genetic factor and the shared family environmental factor contributing to each first-order/second-order cognitive ability associated with the metacognition task and the mentalizing task, we used the structural equation models (SEMs) to fit with the variance-covariance matrix between the MZ and DZ twins (Boker et al., 2011; Keller and Coventry, 2005). The differences in covariance identified the SEM parameters by comparing covariance between MZ and DZ twins along the observed variable as follows

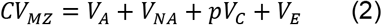

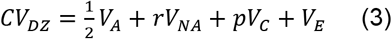

where CV_MZ_ is the covariance between the MZ twins, CV_DZ_ is the covariance between the DZ twins, V_A_ is the additive variance, and *r* is a coefficient for the non-additive variance (V_NA_), which is set as 1/4 or 0, p is a binary coefficient (0 or 1) for the common environmental variance (V_C_), and V_E_ is the error variance including unshared environmental variance. If the MZ resemblance is larger than twice the DZ resemblance, then V_NA_ (or dominant genetic effect, D) is considered, but V_c_ is then ignored. That is *r* is 1/4 and p is 0, and this model is referred to as the ADE model. Otherwise, *r* is 0 and p is 1, and this model is referred to as the ACE model.

To identify which model that should best account for each behavioral measure, we used two model-selection statistics. The first was the chi-square goodness-of-fit statistic. Large values indicate poor model fit to the observed covariance. When two models are nested (i.e., identical except for constraints placed on the sub-model), the difference in fittings between them can be evaluated with the chi-square difference (Δχ^2^). When the chi-square difference is not statistically significant, the more parsimonious model is selected, as the test indicates that the constrained model fits equally well with the data. The second model-selection statistic was the Akaike information criterion (AIC) calculated as follows

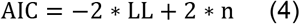

where LL is the log-likelihood of the model on the estimated dataset, and n is the number of parameters in the model. We analyzed the data using maximum-likelihood (ML) methods implemented by the OpenMx module (version: 2.12.2, Boker et al., 2011) in R (version: 3.5.3).

However, it is very likely that the common environmental, additive and non-additive genetic factors simultaneously affect the measured phenotypes (Keller and Coventry, 2005). Thereby, for instance, ignorance of V_NA_ in the ACE model should cause V_C_ underestimated. To correct the potential biases in the conventional ACE and ADE models, we collapsed the additive and non-additive genetic factors to a single genetic factor (V_G_ = V_A_ + V_NA_), and the SEM model is modified as a *q*-GCE model as follows,

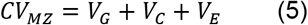

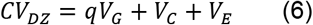

where

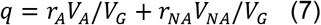

represents the ratio of the genetic factors in the DZ twins in reference to the MZ twins, *r_A_* and *r_NA_* are the ratios in the additive and non-additive genetic factors, respectively. For instance, the assortative mating effect (*i.e*., similar mates are likely married) that would cause overestimates of the common environmental factor can be absorbed into the genetic factor with a value of *r_A_* larger than 1/2. In the conventional ACE model, the value of *q* is 1/2. However, by taking account of the other non-genetic effects other than the shared family environmental effect, the value of *q* might be larger than 1/2. To systematically evaluate the reliability and tolerance of the genetic or shared family environmental effects on each behavioral measure with different values of *q*, we compared the genetic and shared family environmental contributions by continuously changing the value of *q* from 0.05 to 0.95 (**Figure 4**).

#### Bootstrapping

To examine the reliability of the behavioral measures from the empirical data, we repeated 100,000 times of randomly sampling three quarters of all the MZ and DZ twins to measure the Pearson’s correlation and Cronbach’s alpha on the test-retest observations in the metacognition task. Further, to examine the reliability of familial resemblance, we also repeated 100,000 times of randomly sampling three quarters of all the MZ and DZ twins to calculate the ICCs in MZ and DZ twins using the similar bootstrap procedure. Finally, to examine the reliability of the selected model and the model parameters, we also repeated 100,000 times of randomly sampling three quarters of all the MZ and DZ twins to compare the SEM models best fitting with the observations using the similar bootstrap procedure.

#### Power analyses

We made post-hoc power analyses to assess how often the conclusions from the empirical data could be drawn if the same covariance matrices of the MZ and DZ twins are repeatedly obtained by randomly sampling the twins with the same sample size. To do so, we used the OpenMx module (version: 2.12.2) in R (version: 3.5.3) specific for power analysis in the twin study (Boker et al., 2011; Neale and Cardon, 1992). In particular, we examined the selectivity of contribution 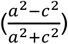 between the genetic factor and the shared family environmental factor in each simulation (**Figure 4 – figure supplement 1B**). Further, we also estimated the expected power at the different sample size with the same covariance matrices of the MZ and DZ twins (**Figure 4 – figure supplement 1A**).

## Author Contributions

S. Jiang and F. Sun conducted the experiments; S. Jiang and X. Wan analyzed the data; P. Yuan assisted with the data collection; Y. Jiang and X. Wan designed the experiments, wrote the manuscript and supervised the project.

## Competing Interest Statement

There were no conflict interests.

## Acknowledgments

This research was funded by the National Natural Science Foundation of China (No. 31471068, X.W. and No. 31830037, Y.J.), the Key Program for International S&T Cooperation Projects of China (MOST, 2016YFE0129100, X.W.), the Fundamental Research Funds for the Central Universities (2017EYT33, X.W.), and also supported by BeTwiSt of Institute of Psychology, Chinese Academy of Sciences.

## Data Availability

The data and source codes are deposited in https://github.com/ShaohanJiang/originsMetacognition.git.

---

**Figure 1 - figure supplement 1.**
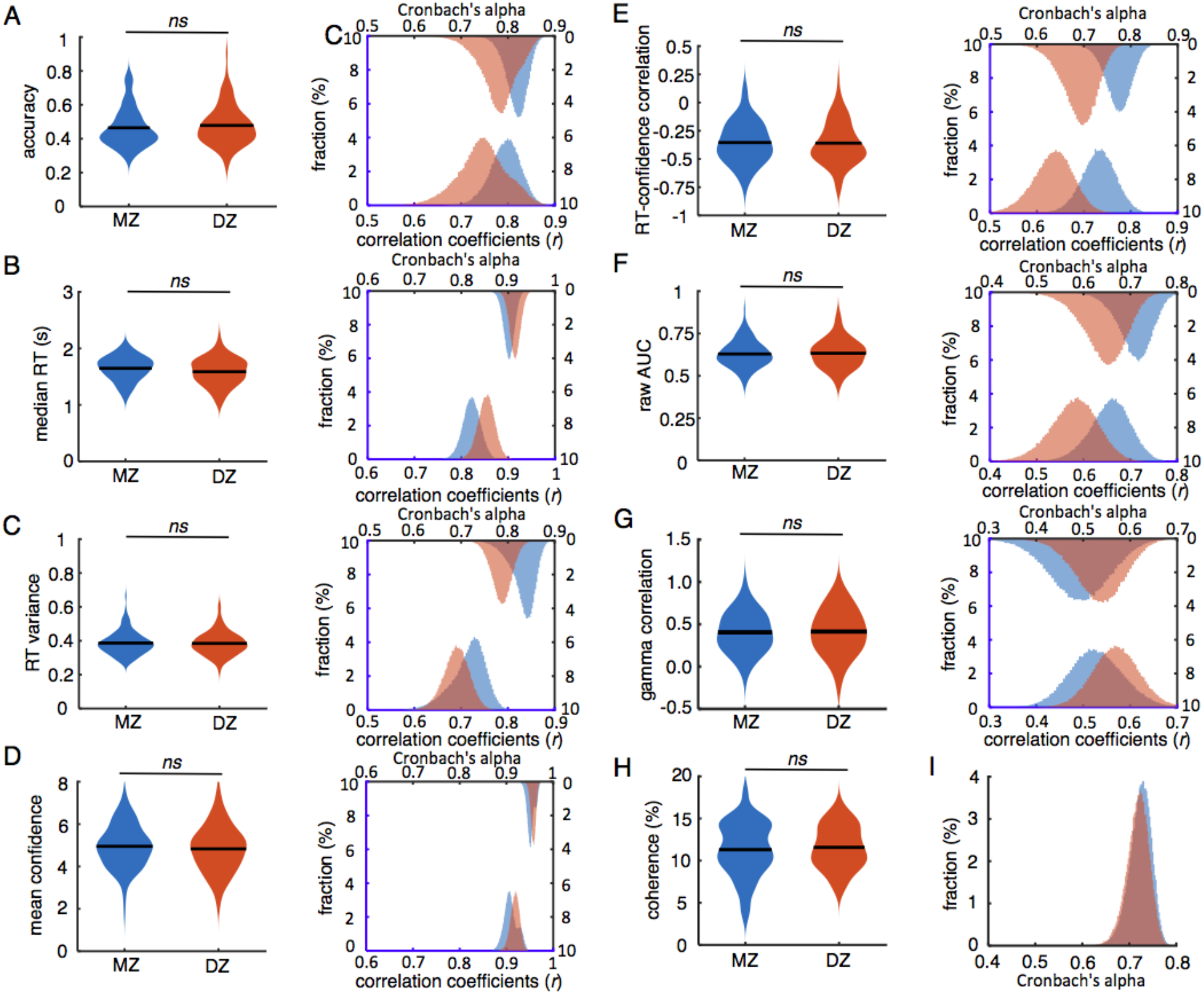
The MZ and DZ twins’ behavioral measures and the reliabilities of these behavioral measures in the metacognition task. (**A**) The accuracy; the mean and population (left); the Pearson’s correlation coefficients and the Cronbach’s alpha between the two repeated metacognition tasks using the bootstrapping procedure (right). (**B**) The median RT. (**C**) The RT variance. (**D**) The mean confidence. (**E**) The RT-confidence correlation. (**F**) The raw AUC. (**G**) The gamma correlation between the confidence and the correctness. (**H**) The coherence that was always kept constant across the two repeated metacognition tasks for each participant. (**I**) The Cronbach’s alpha across the three second-order behavioral measures of the metacognitive abilities. *ns*, no significance.

**Figure 1 - figure supplement 2.**
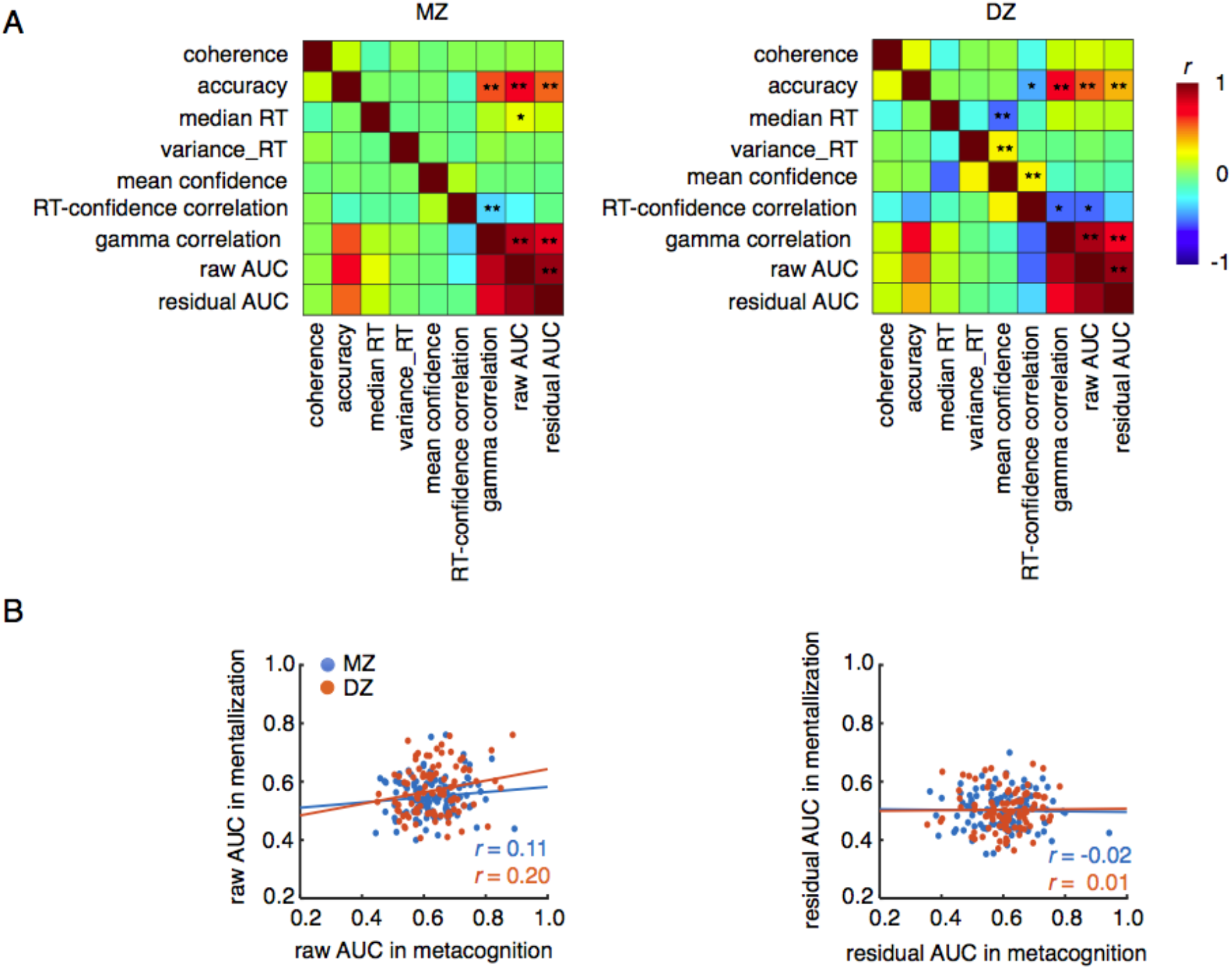
Correlations among the behavioral measures in the metacognition task and between the metacognition and mentalizing tasks. (**A**) Mutual correlations among the behavioral measures in the metacognition task. (**B**) The correlation between the raw AUCs (left) and residual AUCs (right) in the metacognition task and the mentalizing task, respectively. \**P* < 0.05; \*\**P* < 0.01; \*\*\**P* < 0.001.

**Figure 2 - figure supplement 1.**
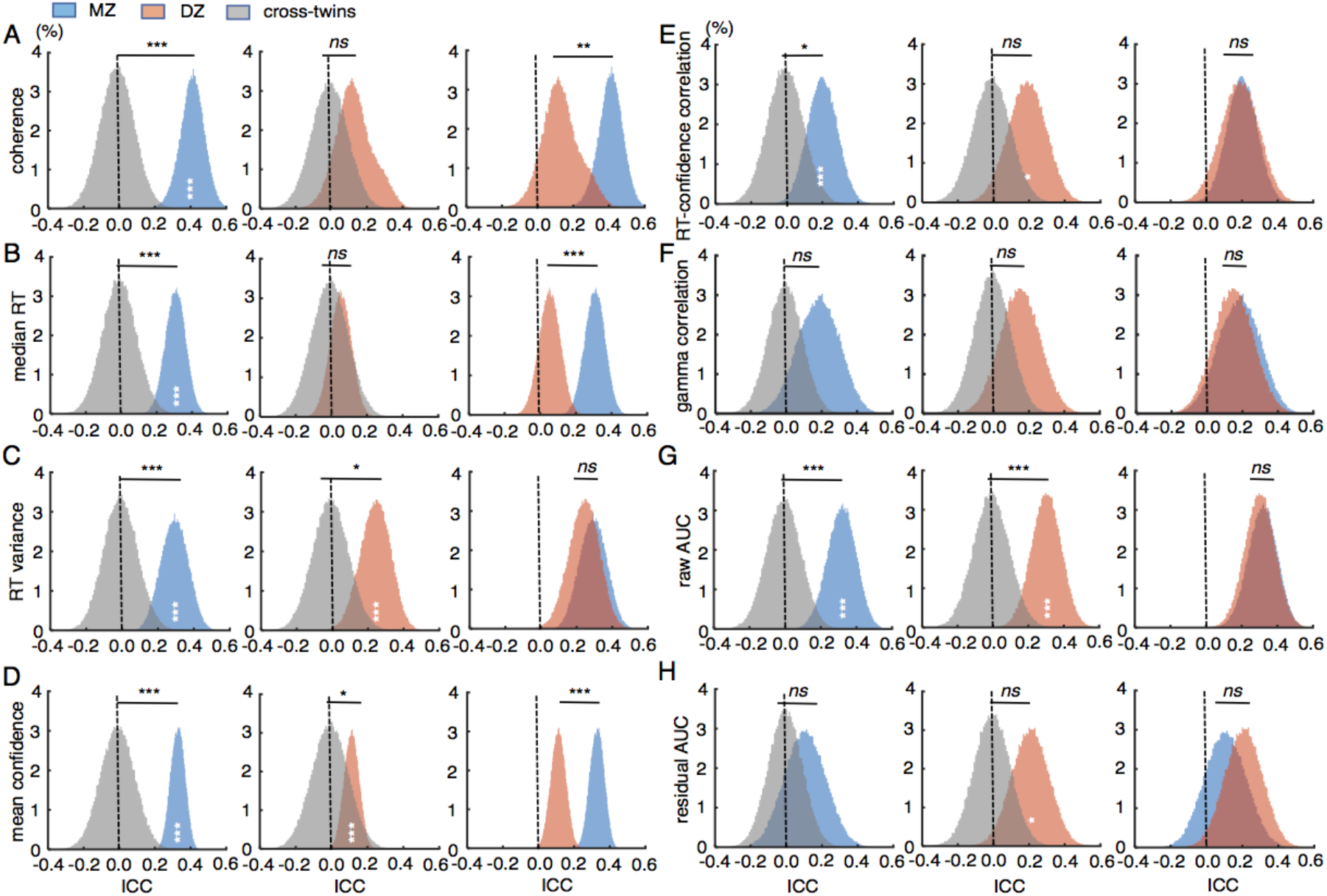
The ICCs of the behavioral measures in the metacognition task using the bootstrapping procedure. (**A**) The within-twin ICCs of the coherence in the MZ and DZ twins and the cross-twin ICCs between the two groups. (**B**) The ICCs of the median RT; (**C**) The ICCs of the RT variance. (**D**) The ICCs of the mean confidence. (**E**) The ICCs of the RT-confidence correlation; (**F**) The ICCs of the gamma correlation between the confidence and the correctness. (**G**) The ICCs of the raw AUC. (**H**) The ICCs of the residual AUC after the RT-associated component was regressed out. The upper statistic sign indicate the significance of the two populations in each figure, and that within each population indicates the significance of the population from zero (dotting black lines). *ns*, no significance; \**P* < 0.05; \*\**P* < 0.01; \*\*\**P* < 0.001.

**Figure 3 - figure supplement 1.**
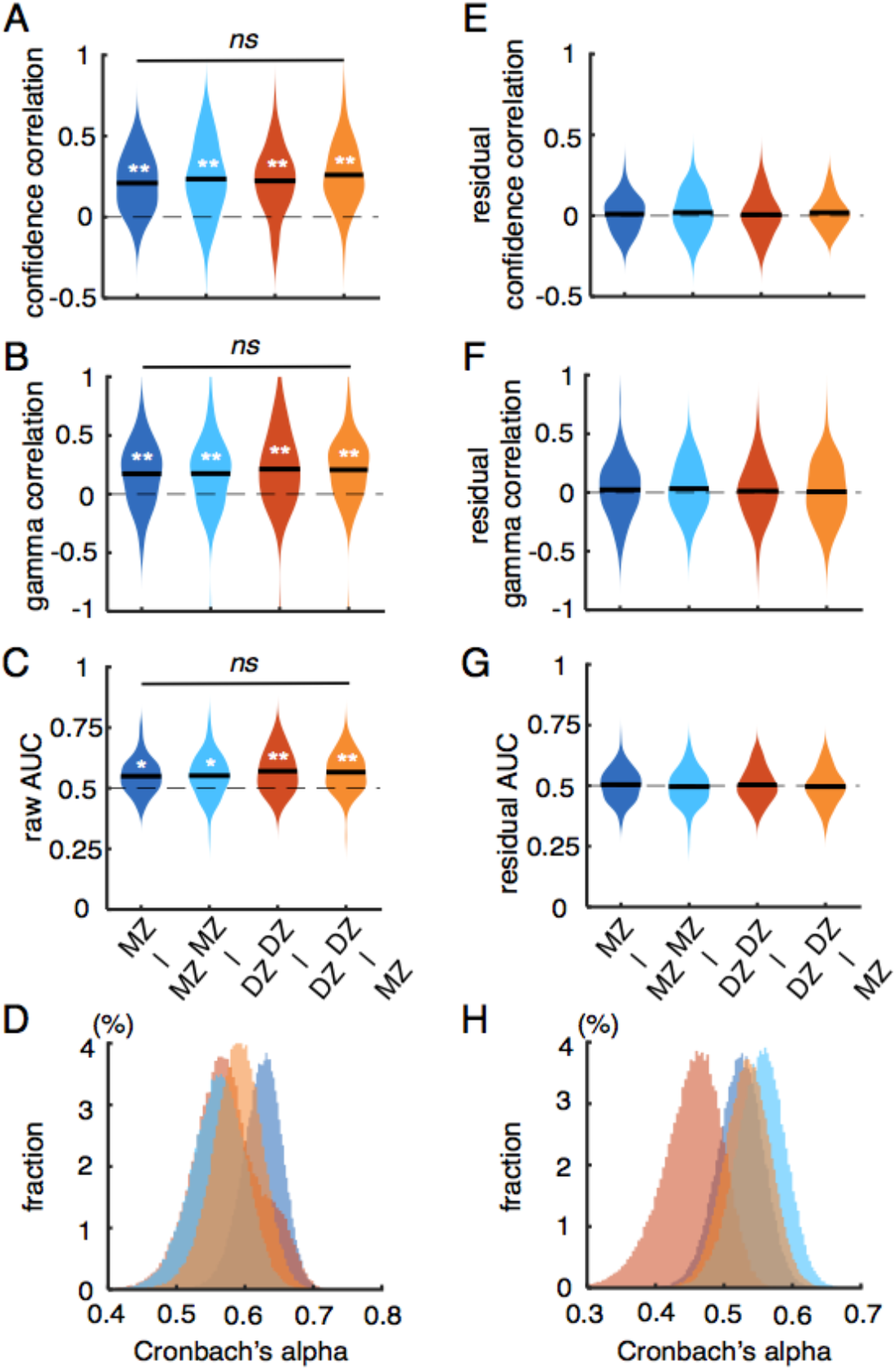
The MZ and DZ twins’ behavioral measures and the consistence of these behavioral measures in the mentalizing task. (**A**) Correlation between the estimated confidence and the other’s reported confidence. (**B**) Gamma correlation between the estimated confidence and the other’s correctness. (**C**) Raw AUC measuring the estimated confidence and the other’s correctness. (**D**) Cronbach’s alpha across the three behavioral measures in each condition. (**E**) Correlation between the residuals of estimated confidence and the other’s reported confidence after the RT-associated component was regresses out. (**F**) Gamma correlation between the residuals of estimated confidence and the other’s correctness after the RT-associated component was regresses out. (**G**) Residual AUC after the RT-associated component was regresses out. (**H**) Cronbach’s alpha across the three behavioral measures after the RT-associated component was regresses out in each condition. MZ-MZ, The MZ participant estimated the partner from the same MZ twin; MZ-DZ, The MZ participant estimated the partner from the DZ twin; DZ-DZ, The DZ participant estimated the partner from the same DZ twin; DZ-MZ, The DZ participant estimated the partner from the MZ twin. \**P* < 0.05; \*\**P* < 0.01 after Bernoulli correction.

**Figure 3 - figure supplement 2.**
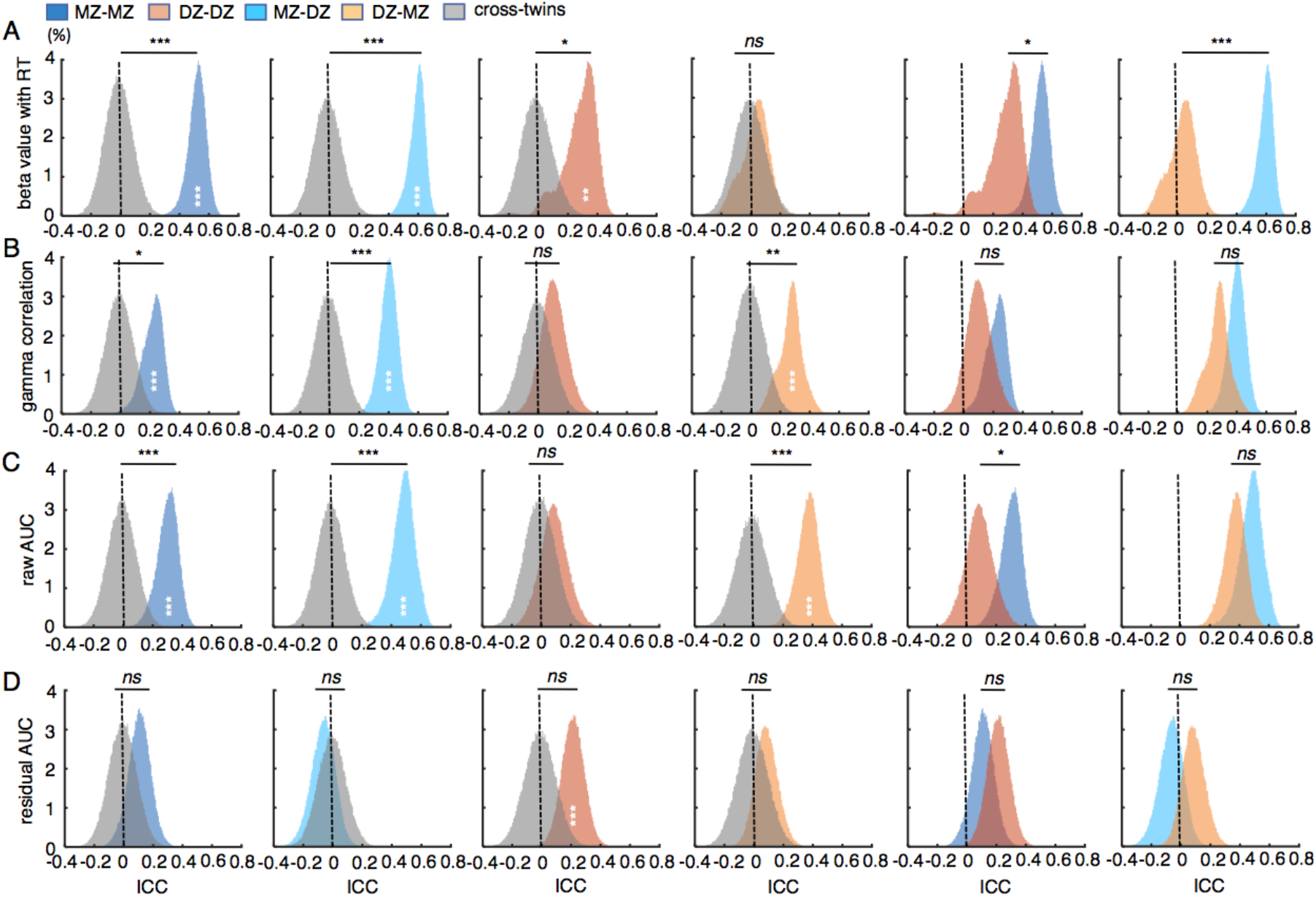
The ICCs of the behavioral measures in the mentalizing task using the bootstrapping procedure. (**A**) The ICCs of the beta values of the regression between the estimated confidence and the other’s RTs in the four within-twin conditions and the cross-twins condition. (**B**) The ICCs of gamma correlation between the estimated confidence and the other’s correctness; (**C**) The ICCs of raw AUC. (**D**) The ICCs of the residual AUC after the RT-associated component was regressed out. The upper statistic sign indicate the significance of the two populations in each figure, and that within each population indicates the significance of the population from zero (dotting black lines). *ns*, no significance; \**P* < 0.05; \*\**P* < 0.01; \*\*\**P* < 0.001.

**Figure 4 - figure supplement 1.**
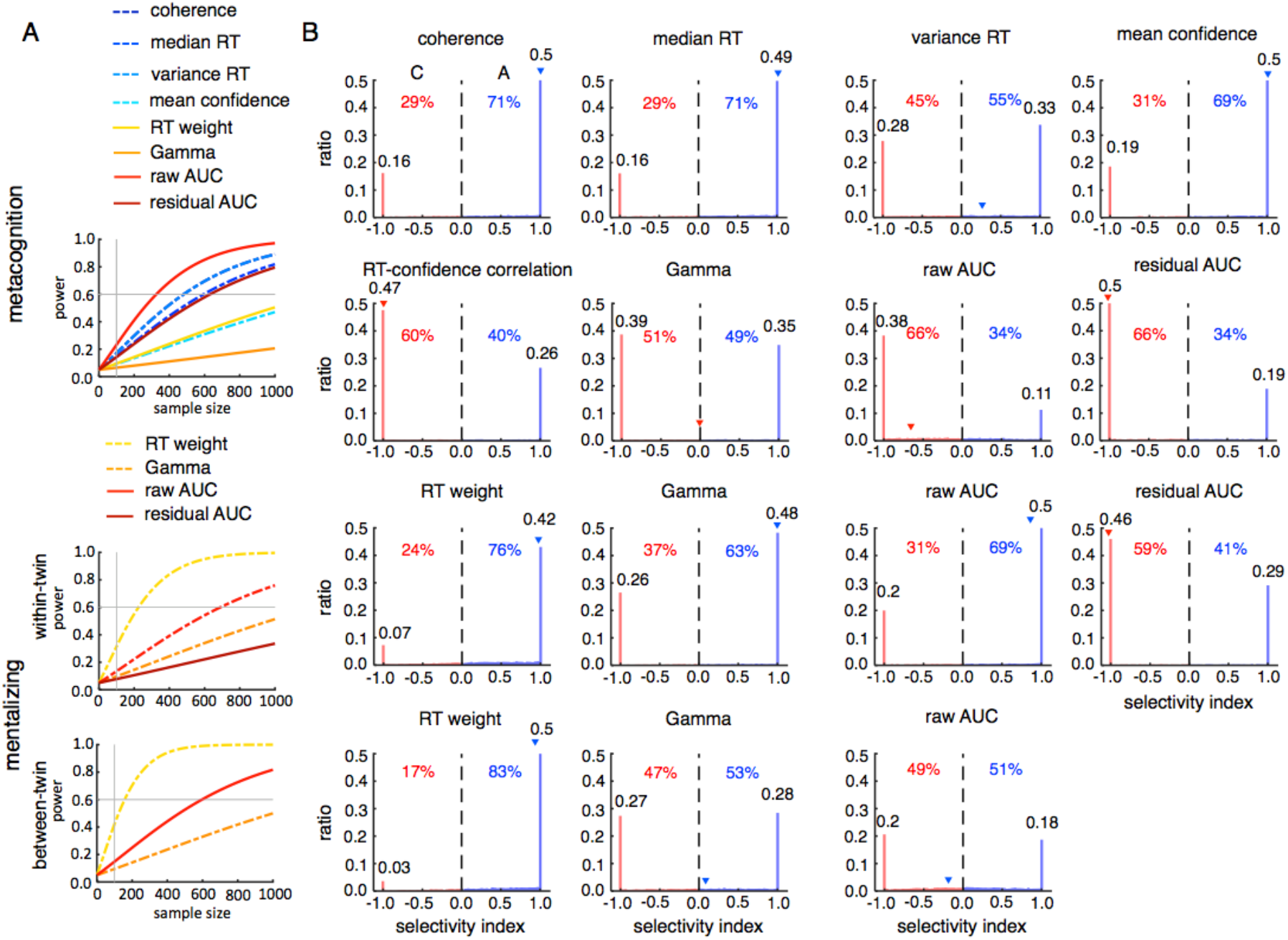
The post-hoc power analyses. (**A**) The power changed with the sample size including both the MZ and DZ twins. The vertical gray lines indicate the same sample size as used in the present study. (**B**) The selectivity of contribution between the genetic factor and the shard family environmental factor in each of simulation with the same sample size as used in the present study. The left side represents biases to the shared family environmental factor, and the right side represents biases to the genetic factor. The percentage numerical values indicate the total percentage over the left and right sides, respectively, and the decimal values indicate the ratio at the selectivity index of −1 and 1, respectively. The triangle marker indicates the selectivity index of the empirical data.

## Notes

### Competing Interest Statement

The authors have declared no competing interest.

## References

Atzil S, Gao W, Fradkin I, Barrett LF (2018) Growing a social brain Nature Human Behaviour 2: 624–636.

Baker L (1994) Fostering metacognitive development Advances in child development and behavior 25: 201–239.

Boker S, Neale M, Maes H, Wilde M, Spiegel M, Brick T, Spies J, Estabrook R, Kenny S, Bates T (2011) OpenMx: an open source extended structural equation modeling framework Psychometrika 76: 306–317.

Colclough GL, Smith SM, Nichols TE, Winkler AM, Sotiropoulos SN, Glasser MF, Van Essen DC, Woolrich MW (2017) The heritability of multi-modal connectivity in human brain activity Elife 6: e20178. doi:10.7554/eLife.20178

Davies G, Marioni RE, Liewald DC, Hill WD, Hagenaars SP, Harris SE, Ritchie SJ, Luciano M, Fawns-Ritchie C, Lyall D (2016) Genome-wide association study of cognitive functions and educational attainment in UK Biobank (N= 112 151) Molecular psychiatry 21: 758–767.

Deary IJ, Weiss A, Batty GD (2010) Intelligence and personality as predictors of illness and death: How researchers in differential psychology and chronic disease epidemiology are collaborating to understand and address health inequalities Psychological science in the public interest 11: 53–79.

Feurer E, Sassu R, Cimeli P, Roebers C (2015) Development of meta-representations: Procedural metacognition and the relationship to theory of mind Journal of Educational and Developmental Psychology 5: 6–18.

Flavell JH (1979) Metacognition and cognitive monitoring: A new area of cognitive–developmental inquiry American psychologist 34: 906.

Flavell JH (1999) Cognitive development: Children’s knowledge about the mind Annual review of psychology 50: 21–45.

Flavell JH (2003) Varieties of uncertainty monitoring Behavioral and Brain Sciences 26: 344.

Fleming SM, Weil RS, Nagy Z, Dolan RJ, Rees G (2010) Relating introspective accuracy to individual differences in brain structure Science 329: 1541–1543.

Fleming SM, Huijgen J, Dolan RJ (2012). Prefrontal contributions to metacognition in perceptual decision-making. J. Neurosci. 32: 6117–6125.

Frith CD, Frith U (2012) Mechanisms of social cognition Annual review of psychology 63: 287–313.

Galton F (1870) Hereditary genius: An inquiry into its laws and consequences D. Appleton.

Gao W, Elton A, Zhu H, Alcauter S, Smith JK, Gilmore JH, Lin W (2014) Intersubject variability of and genetic effects on the brain’s functional connectivity during infancy Journal of Neuroscience 34: 11288–11296.

Goupil L, Kouider S (2016) Behavioral and neural indices of metacognitive sensitivity in preverbal infants Current Biology 26: 3038–3045.

Grayson D (1989) Twins reared together: Minimizing shared environmental effects Behavior genetics 19: 593–604.

Greven CU, Harlaar N, Kovas Y, Chamorro-Premuzic T, Plomin R (2009) More than just IQ: School achievement is predicted by self-perceived abilities—But for genetic rather than environmental reasons Psychological Science 20: 753–762.

Heyes C, Bang D, Shea N, Frith CD, Fleming SM (2020) Knowing ourselves together: the cultural origins of metacognition Trends in cognitive sciences 24: 349–362.

Heyes CM, Frith CD (2014) The cultural evolution of mind reading Science 344.

Hill J, Inder T, Neil J, Dierker D, Harwell J, Van Essen D (2010) Similar patterns of cortical expansion during human development and evolution Proceedings of the National Academy of Sciences 107: 13135–13140.

Hill WG, Goddard ME, Visscher PM (2008) Data and theory point to mainly additive genetic variance for complex traits PLoS Genet 4: e1000008.

Hughes C, Jaffee SR, Happé F, Taylor A, Caspi A, Moffitt TE (2005) Origins of individual differences in theory of mind: From nature to nurture? Child development 76: 356–370.

Jansen AG, Mous SE, White T, Posthuma D, Polderman TJ (2015) What twin studies tell us about the heritability of brain development, morphology, and function: a review Neuropsychology review 25: 27–46.

Jiang S, Wang S, Wan X. Distinct mental state representations of decision uncertainty in mentalizing and metacognition. (submitted).

Johnson W, Turkheimer E, Gottesman II, Bouchard Jr TJ (2009) Beyond heritability: Twin studies in behavioral research Current directions in psychological science 18: 217–220.

Kaas JH (2006) Evolution of the neocortex Current Biology 16: R910–R914.

Kahneman D, Lovallo D (1993) Timid choices and bold forecasts: A cognitive perspective on risk taking Management science 39: 17–31.

Keller MC, Coventry WL (2005) Quantifying and addressing parameter indeterminacy in the classical twin design Twin Research and Human Genetics 8: 201–213.

Kiani R, Shadlen MN (2009) Representation of confidence associated with a decision by neurons in the parietal cortex Science 324: 759–764.

Kliemann D, Adolphs R (2018) The social neuroscience of mentalizing: challenges and recommendations Current opinion in psychology 24: 1–6.

Knafo A, Uzefovsky F (2013) Variation in empathy: The interplay of genetic and environmental factors.

Kruger J, Dunning D (1999) Unskilled and unaware of it: how difficulties in recognizing one’s own incompetence lead to inflated self-assessments Journal of personality and social psychology 77: 1121.

Kunimoto C, Miller J, Pashler H (2001). Confidence and auucary of near-threshold discrimination repsonses. Conscious. Cogn. 10, 294–340.

Levitt H (1971) Transformed up - down methods in psychoacoustics The Journal of the Acoustical society of America 49: 467–477.

Lurz RW (2011) Mindreading animals: The debate over what animals know about other minds MIT press.

McGraw KO, Wong SP (1996) Forming inferences about some intraclass correlation coefficients Psychological methods 1: 30.

Mueller S, Wang D, Fox MD, Yeo BT, Sepulcre J, Sabuncu MR, Shafee R, Lu J, Liu H (2013) Individual variability in functional connectivity architecture of the human brain Neuron 77: 586–595.

Neale M, Cardon LR (1992) Methodology for genetic studies of twins and families (Vol. 67) Springer Science& Business Media.

Nelson TO (1990) Metamemory: A theoretical framework and new findings In Psychology of learning and motivation (Vol. 26: pp. 125–173) Elsevier.

Onishi KH, Baillargeon R (2005) Do 15-month-old infants understand false beliefs? Science 308: 255–258.

Plomin R, Daniels D (2011) Why are children in the same family so different from one another? Int J Epidemiol 40: 563–582. doi:10.1093/ije/dyq148

Plomin R, Deary IJ (2015) Genetics and intelligence differences: five special findings Molecular psychiatry 20: 98–108.

Plomin R, Scheier MF, Bergeman CS, Pedersen NL, Nesselroade JR, McClearn GE (1992) Optimism, pessimism and mental health: A twin/adoption analysis Personality and individual differences 13: 921–930.

Polderman TJ, Benyamin B, De Leeuw CA, Sullivan PF, Van Bochoven A, Visscher PM, Posthuma D (2015) Meta-analysis of the heritability of human traits based on fifty years of twin studies Nature genetics 47: 702–709.

Qiu L, Su J, Ni Y, Bai Y, Zhang X, Li X, Wan X (2018) The neural system of metacognition accompanying decision-making in the prefrontal cortex PLoS biology 16: e2004037.

Sternberg RJ (1984) Toward a triarchic theory of human intelligence Behavioral and Brain Sciences 7: 269–287.

Strenze T (2007) Intelligence and socioeconomic success: A meta-analytic review of longitudinal research Intelligence 35: 401–426.

Taumoepeau M, Ruffman T (2008) Stepping stones to others’ minds: Maternal talk relates to child mental state language and emotion understanding at 15, 24, and 33 months Child development 79: 284–302.

Tucker-Drob EM, Briley DA, Harden KP (2013) Genetic and environmental influences on cognition across development and context Current directions in psychological science 22: 349–355.

Turkheimer E (2000) Three laws of behavior genetics and what they mean Current directions in psychological science 9: 160–164.

Vygotsky L (1978) Mind in society Cambridge, MA: MIT Press.

Warrier V, Grasby KL, Uzefovsky F, Toro R, Smith P, Chakrabarti B, Khadake J, Mawbey-Adamson E, Litterman N, Hottenga J-J (2018) Genome-wide meta-analysis of cognitive empathy: heritability, and correlates with sex, neuropsychiatric conditions and cognition Molecular psychiatry 23: 1402–1409.

